# Alpha and Beta Oscillations Mediate the Effect of Motivation on Neural Coding of Cognitive Flexibility

**DOI:** 10.1101/2025.03.11.642697

**Authors:** Juan M. Chau, Matias J. Ison, Paul S. Muhle-Karbe, Mark G. Stokes, Sam Hall-McMaster, Nicholas E. Myers

**Author notes:** Joint last authors (Sam Hall-McMaster, Nicholas E. Myers).

## Abstract

Cognitive flexibility is crucial for adaptive human behaviour. Prior studies have analysed the effect of reward on cognitive flexibility; however, the neural mechanisms underlying these effects remain largely unknown. This study explores how reward influences neural oscillations and how these changes impact behavioural performance. Using time-frequency decomposition, we examined electroencephalographic data from participants engaged in rule-guided task-switching with varying reward prospects. Higher anticipated rewards lead to greater desynchronisation of alpha (8-12Hz) and beta (20-30Hz) oscillations, which in turn correlated with improved task performance. Both alpha power and event-related potential (ERP) coding of reward independently predicted reward-based performance improvements, suggesting distinct mechanisms supporting proactive control. These findings underscore the unique contributions of neural oscillations in mediating motivational effects on cognitive flexibility.

## Introduction

Understanding the neural mechanisms underlying cognitive flexibility and motivation is crucial for advancing our understanding of how these processes support adaptive behaviour in complex environments. For instance, in everyday life, cognitive flexibility allows individuals to switch between different tasks efficiently, such as a parent juggling work responsibilities while helping their child with homework. Motivation plays a key role in this process, as a motivated individual is more likely to stay focused and adapt to changing demands (Braem & Egner, 2018). Similarly, in sports, an athlete’s ability to adjust their strategy based on the opponent’s moves and stay motivated, especially throughout high-stakes events, can significantly impact their performance (Fletcher & Sarkar, 2012).

Cognitive flexibility and the related concept of cognitive control encompass both reactive and proactive control mechanisms (Braver et al., 2009; Braver, 2012). Reactive control involves spontaneous adjustment of behaviour in response to unexpected changes in task demands, often relying on immediate cues to guide actions. In contrast, proactive control entails the anticipatory maintenance of task-relevant information in situations where fast and accurate responses are incentivized. This latter type of control has been linked to midfrontal neural oscillations, particularly in the theta band (4-8 Hz; Cooper et al., 2016; Kaiser & Schütz-Bosbach, 2019). These oscillations are thought to play a role in preparatory adjustments that optimize performance in response to expected task demands. Proactive control is essential for cognitive flexibility, allowing individuals to adapt their behaviour based on anticipation of changing task requirements and reward contingencies. However, how motivation influences these mechanisms is still poorly understood.

Hall-McMaster et al. (2019) investigated which aspects of sequential task processing are affected by reward to optimise cognitive flexibility. They found that high reward prospects increased neural representations for task rule information, which was associated with performance improvements. However, they raised an open-ended question about the exact mechanism by which a stronger separation of task representations translates into performance improvements. This question suggests that while reward enhances the clarity and distinctiveness of neural coding, the neural mechanisms through which motivation influences overall cognitive and behavioural performance remain to be fully understood.

One possibility comes from sharpening task control through oscillations in the alpha band (8-12 Hz). Alpha oscillations have been implicated in various cognitive functions, including attention and memory (Klimesch, 2012; Palva & Palva, 2007; Rihs et al., 2009). For instance, desynchronisation of alpha oscillations may contribute to the disinhibition of task-relevant areas to facilitate anticipated stimulus processing (Gould et al., 2011; Haegens et al., 2011; Worden et al., 2000). Prior studies have shown that alpha power can be modulated by task demands and reward, influencing behavioural outcomes such as response times (RTs; Pessoa & Engelmann, 2010; Van Dijk et al., 2008). For instance, a decrease in alpha power has been associated with enhanced cognitive control and faster RTs in high-reward conditions, suggesting a link between motivation and neural efficiency (Arnau et al., 2024; Hughes et al., 2013; van Driel et al., 2015). Additionally, alpha oscillations have been found to play a role in the suppression of irrelevant (Foxe & Snyder, 2011; Jensen & Mazaheri, 2010; Klimesch, 1999) or less-rewarding (Heuer et al., 2017) information, thereby facilitating task performance. While oscillations have been used to study various aspects of cognitive control, they have not been extensively explored in the context of information coding and cognitive flexibility. Our prediction is that the effect of reward coding on cognitive flexibility is mediated by alpha oscillations.

Oscillatory responses in the alpha band may serve as neural markers of how effectively individuals translate motivational incentives into cognitive performance (Knyazev, 2007; Messerotti Benvenuti et al., 2019; Zhu et al., 2019). Examining individual variability in these oscillatory dynamics provides a valuable opportunity to understand how reward prospect interacts with cognitive flexibility at the neural level. This is particularly relevant given that motivational states can modulate attentional allocation and task preparation, with variability across individuals reflecting differences in neural responsiveness to reward cues (Braem et al., 2012; Lee & Reeve, 2017). Reward sensitivity could therefore play a key role in shaping how individuals engage with cognitive control demands.

In this study, we aimed to investigate how reward anticipation modulates neural oscillations, and how these changes relate to cognitive flexibility. We hypothesised that higher anticipated reward would lead to greater desynchronisation in the alpha band, reflecting enhanced attentional engagement and proactive control. We also explored whether oscillatory changes in other frequency bands (e.g., beta oscillations) contributed to this effect. Finally, we examined whether these oscillatory dynamics as well as the multivariate neural coding of reward made independent contributions to behaviour to better understand how distinct neural processes support motivated cognitive control.

## Methods

### Data source

The data for this study were from a previously published experiment investigating the impact of reward on neural coding of task rules. The original dataset was previously described in Hall-McMaster et al. (2019) and is publicly available at https://osf.io/kuzye/.

### Participants

Thirty participants aged between 18 and 35 years (mean age: 23 years; 19 females) took part in the study. A post hoc sensitivity analysis using the *pwr* R package (version 1.3-0; Champely, S., 2020) indicated that this sample size provides 80% power to detect medium-sized correlations (e.g., ρ≥0.4866) between oscillatory changes and behavioural measures. All participants had normal or corrected-to-normal vision and reported no history of neurological or psychiatric conditions. Participants were compensated at a rate of £8 per hour or received course credit, with the potential to earn up to an additional £10 based on their performance. Ethical approval was granted by the Central University Research Ethics Committee at the University of Oxford, and all participants provided informed consent prior to participation.

### Experimental setup and stimuli

The original data collection involved presenting stimuli on a 22-inch screen with a spatial resolution of 1280 × 1024 and a refresh rate of 60 Hz. Stimulus presentation was controlled using Psychophysics Toolbox-3 (Kleiner et al., 2007). Reward cues and feedback were displayed in size 30 Arial font. Task cues and target stimuli had visual angles of approximately 2.52° (100 × 100 pixels) and 1.26° (50 × 50 pixels), respectively, based on an estimated viewing distance of 60 cm. Responses were recorded using the F and J keys on a standard QWERTY keyboard.

### Experimental design

In this task (Fig. 1), participants aimed to accumulate as many points as possible by categorizing bidimensional target stimuli based on either their colour (yellow vs. blue) or shape (square vs. circle). On each trial one feature dimension (colour or shape) was task-relevant while the other served as an irrelevant distractor. The relevant feature dimension was indicated by a visual task cue presented before the target onset. Additionally, each trial offered either a high or low reward magnitude for correct responses, signalled at the beginning of each trial by a single pound sign (£; low reward, 5–10 points) or three pound signs (£££; high reward, 50–100 points).

**Figure 1.**
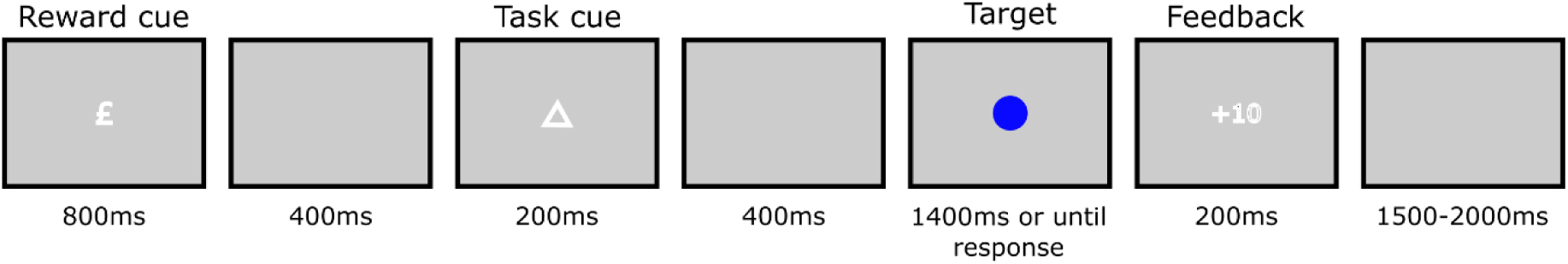
Experimental design. On each trial, the reward cue was presented, followed by an empty-screen delay. A task cue was presented, indicating the relevant feature of the upcoming stimulus to respond to. After a second empty-screen delay, a coloured shape appeared until the participant responded, for a maximum time of 1.4 seconds. Feedback based on both accuracy and speed were given before the inter-trial empty-screen delay.

On each trial, the reward cue (£ or £££) was displayed for 800 ms, followed by a 400 ms delay. Next, the task cue (one of four abstract shapes) appeared for 200 ms, with the mapping of two cues to each task counterbalanced between participants. After a 400 ms delay, the target (a yellow square, blue square, yellow circle, or blue circle) was presented and remained on screen until a response was made or for a maximum of 1400 ms. If the cued task rule was colour, the correct response mapping was *F* for yellow and *J* for blue. If the rule was shape, *F* corresponded to square and *J* to circle. The response phase was followed by 200 ms of feedback. Incorrect responses or omissions resulted in feedback showing 0 points, while correct responses displayed X points, where X varied within the high or low reward ranges based on response time (RT).

We aimed to incentivise fast responses by dynamically adapting the reward level to each participant’s response time distribution. First the RT threshold for different points was initialized so that responses faster than 400, 600, 800, 1000, 1200, and 1400 ms earned 100 (10), 90 (9), 80 (8), 70 (7), 60 (6), and 50 (5) points on high (low) reward trials, respectively. We then built up a participant’s RT distribution. For correct trials, the current trial RT was added to an array for its reward condition. Once each reward level contained more than six values, individualized points criteria were recalculated following Shen and Chun (2011). Responses faster than 95%, 80%, 65%, 50%, and 35% of previous RTs from the same condition were rewarded with the most to least points. For example, if a participant had the following RTs on the first six high-reward trials: 830, 860, 900, 930, 960, and 1000 ms, the thresholds for the seventh high-reward trial would be calculated based on the percentiles of this distribution. A response faster than 95% of the values (e.g., <830 ms) would earn 100 points, while a response faster than 80% of the values (e.g., <860 ms) would earn 90 points. The trial concluded with a randomly selected ITI duration, drawn from a uniform distribution of 1000, 1100, 1200, 1300, or 1400 ms.

Participants were trained to achieve a minimum of 70% accuracy before completing 10 experimental blocks of 65 trials each. Excluding the first trial in each block, equal numbers of reward cues, task cues, stimuli, and ITI durations were presented. The presentation was pseudorandomized to ensure trials were balanced based on task, target congruency, and task sequence for each reward condition. Target congruency referred to whether task-relevant and task-irrelevant features were mapped to the same (congruent) or different (incongruent) response hands. Task sequence referred to whether the task rule was different from the previous trial (switch trial) or the same (repeat trial).

### EEG data acquisition and preprocessing

EEG data were recorded using 61 Ag/AgCl sintered electrodes (EasyCap), a NeuroScan SynAmps RT amplifier, and Curry 7 acquisition software (RRID:SCR_009546). The EEG data were preprocessed in EEGLAB (version 14.1.1b; Delorme and Makeig, 2004). EEG analyses were conducted in MATLAB (version 2022b) using the FieldTrip toolbox (revision 3f959be1b; Oostenveld, 2011) along with custom scripts.

EEG data were down-sampled from 1000 Hz to 250 Hz and filtered using a 0.01 Hz high-pass filter and a 40 Hz low-pass filter. For each participant, channels with excessive noise were identified through visual inspection and replaced via interpolation, using a weighted average of the surrounding electrodes. The data were then rereferenced by subtracting the mean activation across all electrodes from each individual electrode at each time point. Data were segmented into epochs ranging from -1000 to 5000 ms relative to the onset of the reward cue. Epochs containing artifacts, such as muscle activity, were rejected based on visual inspection. The data also underwent independent component analysis to remove stereotyped artifacts such as eye blinks, which were removed through visual inspection. On average, 16.6% of the trials (M=107.73, SD=43.77) were removed per participant.

A Laplacian filter was applied to reduce volume conductivity using the spline method with a Legendre polynomial degree of 14. Power spectral density (PSD) was calculated using an 800 ms Hamming window with a 20 ms step, allowing for a frequency range between 1.9531 Hz and 39.0625 Hz, and a frequency resolution of 0.9766 Hz. This resulted in the dataset used for subsequent analyses. Prior to each analysis, data were z-scored over the trial dimension and baseline corrected using a time window of 600 to 300 ms before the onset of the reward cue, or 250 to 50 ms before the onset of the task or target cues. These baseline windows were selected separately for each phase to maximize information separability related to upcoming task events, while minimizing potential carryover effects from preceding stimuli. The shorter baseline windows before the task and target cues were chosen to match the original study (Hall-McMaster, 2019), and the longer baseline window before the reward cue took advantage of the preceding inter-trial interval, which provides a longer segment of EEG activity for a more accurate baseline estimation. Since we did not perform direct comparisons across phases, this approach allowed us to optimize the sensitivity of each RSA analysis without confounding task phase with distance from the baseline window.

## Data analysis

We explored how reward modulated oscillatory power, and the effect this had on response times. For this, we observed whether there was a significant effect of reward cue on power across the frequency spectrum. We next tested whether power differences could explain behavioural differences between high and low reward trials by performing a median split analysis. We divided participants into two groups based on the difference between response times in low-and high-reward trials and then examined the reward effect on oscillatory power in both groups. We also performed a correlation to explore potential contributions of different frequency bands over time towards response times.

We analysed EEG data with representational similarity analysis (RSA; Kriegeskorte et al., 2008) to replicate the multivariate event-related potential (ERP) neural coding estimation performed by Hall-McMaster et al. (2019). Here, we define neural coding as the variation in multivariate EEG patterns that reflect the brain’s representation of specific task-relevant variables, such as reward level, task rule, or stimulus features. We extended this approach into the time-frequency domain to explore whether multivariate patterns of oscillatory power coded for the different task variables. Throughout the paper, we use *broadband* when referring to ERP data, and *time-frequency* when referring to oscillatory data. RSA was chosen over other classification-based methods such as linear discriminant analysis (LDA) or support vector machines (SVM) because it is particularly well-suited for EEG data, where neural representations of multiple variables are often distributed and overlapping (Grootswagers et al., 2017). Unlike classifiers that focus on maximizing class separation and require careful balancing of data points in each condition for binary discrimination, RSA measures the similarity structure between conditions (Haxby et al., 2014; Kriegeskorte & Kievit, 2013). This approach makes it possible to isolate the contributions of multiple task variables to the neural similarity structure. Moreover, it enables the assessment of how external factors, such as reward, alter the neural similarity structure on a continuous measurement scale. By examining the temporal dynamics of frequency-specific neural coding, we aimed to identify distinct periods during which task-relevant information was most prominently represented.

In addition to time-frequency analysis, we conducted a univariate ERP analysis on the broadband EEG data to examine whether reward cues elicited differences in evoked potentials. Specifically, we compared average ERP amplitudes between low-and high-reward trials to identify whether reward-related differences were also evident in the time-frequency domain.

Both of these analyses were performed considering only the trials in which participants responded correctly, across four different regions of interest (ROI): all 61 channels, frontal (AF3, AFz, AF4, F3, F1, Fz, F2, F4), central (C3, C1, Cz, C2, C4), and posterior channels (O1, Oz, O2, PO3, POz, PO4) to explore regional specificity in representing information.

In all EEG analyses, we used one-dimensional (for broadband analyses) or two-dimensional (for time-frequency-based analyses) cluster-based permutation t-tests (Maris & Oostenveld, 2007). Clusters were defined as adjacent time-frequency points that exceeded a significance threshold, with adjacency determined along the temporal and frequency dimensions using MATLAB’s *bwconncomp* function. A two-tailed cluster-forming threshold of α=0.05 was applied. No minimum cluster size was imposed. Instead, we computed the absolute cluster mass and compared it to a null distribution of maximum cluster masses (i.e., the largest cluster per permutation) obtained from 10,000 random sign-flip permutations, where the sign of each participant was independently flipped, to determine corrected p-values.

### Representational Similarity Analysis

Representational similarity analysis (RSA) is a powerful tool for examining how different cognitive processes are encoded in the brain over time (Kriegeskorte et al., 2008). RSA leverages pattern information that would typically be averaged out in univariate analyses, enhancing its sensitivity to distributed neural activity (Kriegeskorte et al., 2006). This method allows researchers to explore the dynamic nature of neural representations and their modulation by external factors such as reward levels (Hall-McMaster et al., 2019; Luyckx et al., 2019). While this method is commonly used for estimating the neural coding of different task variables from ERP data (Hall-McMaster et al., 2019; Wei et al., 2023), it also allows for additional dimensions such as frequency bands (Sommer et al., 2022), or channels (Riddeaux, 2024) to be explored simultaneously. Time-frequency RSA has the potential to be applied in various domains, including visual perception, memory, and decision-making, revealing how neural representations change with task demands and contextual factors (Furl et al., 2017; Kikumoto & Mayr, 2020). We performed this process twice for each participant, once for neural coding analyses of the broadband data and once for the neural coding analysis of the time-frequency data (Fig. 2). The first step was to calculate the average pattern per condition, obtaining a matrix of channels x time points x frequency points x conditions. Then, the Mahalanobis distance (MD) between each condition was calculated for each time-frequency point combination using the difference between the power in those conditions, as well as the channel covariance matrix through Equation 1.

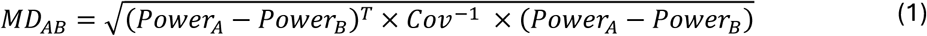

**Figure 2.**
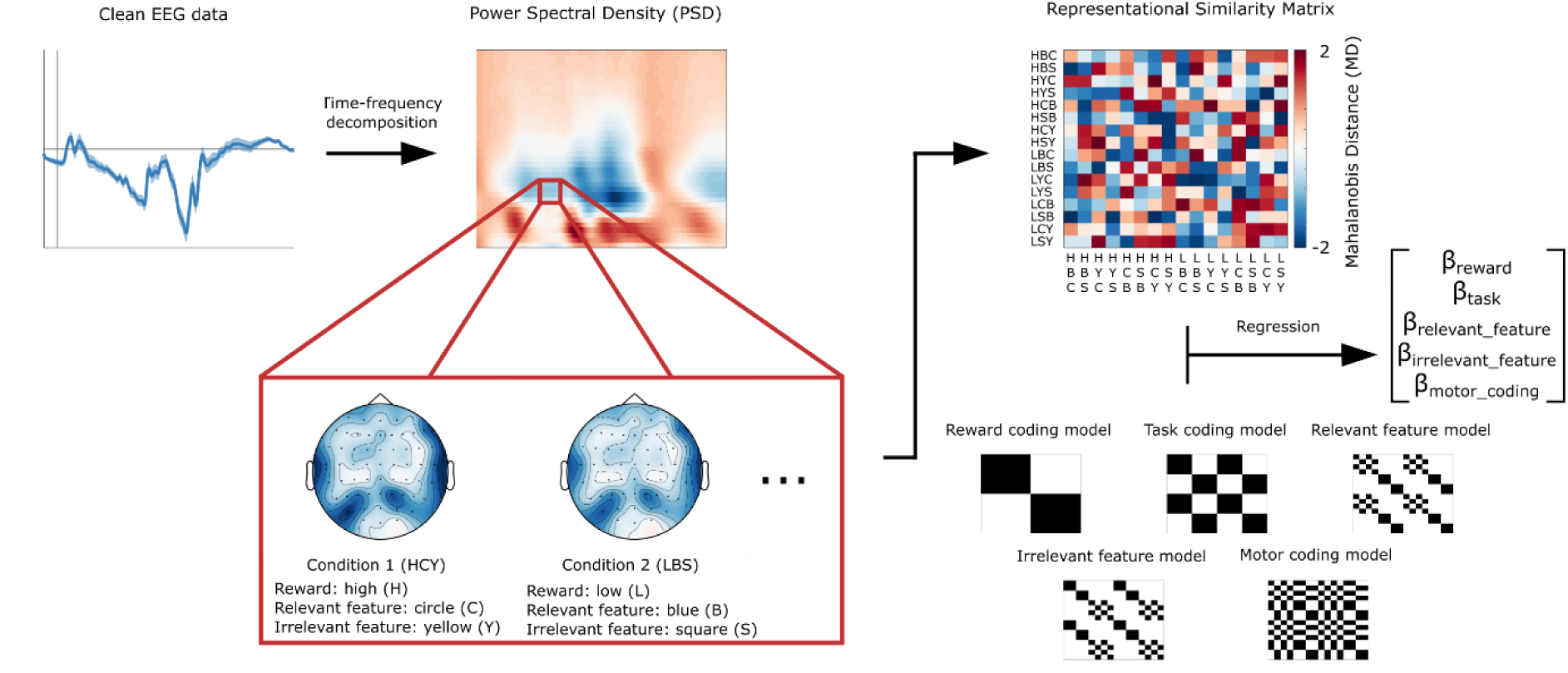
Time-frequency Representational Similarity Analysis (RSA) pipeline. The first step is to calculate the PSD for each EEG channel and trial. Then, Mahalanobis Distances between all 16 conditions averages are calculated for each time and frequency point, resulting in a 16 x 16 dissimilarity matrix. Finally, these dissimilarity matrices are regressed against model RDMs that predict differences in dissimilarity structure for each individual task variable.

We then calculated representational dissimilarity matrices (RDM) of multivariate condition distances for each time-frequency point and for each participant. A total of five 16 x 16 RDMs (two reward levels, two tasks, two colours, and two shapes) were built to capture neural oscillatory patterns related to different task variables (reward, task type, task-relevant and -irrelevant features, and motor coding). These RDMs had zeroes in positions where conditions matched on the variable of interest (e.g., low reward blue square and low reward yellow circle), and ones in the remaining positions. As these RDMs are symmetric, the upper triangular portion of each was transformed into a distance vector (DV) and concatenated before being entered into a linear multiple regression where the time-frequency data (TFD) was the dependent variable, as shown in Equation 2.

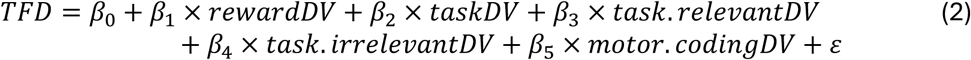

To assess potential multicollinearity among the RDM predictors, we computed variance inflation factors (VIF) for each vectorized RDM. All VIF values were below 2.68, lower than commonly accepted thresholds such as 4 or 10, indicating low multicollinearity (O’Brien, 2007; see Supplementary Table S1). The regression was performed three times considering baseline windows before the onset of the reward, task, and target cues.

## Code availability

The analysis code as well as the preprocessed data can be accessed at https://github.com/JuanMChau/AlphaBetaOscillations.

## Results

### Reward magnitude is encoded in the time-frequency domain along the trial

We used RSA to test for coding of task variables in the time-frequency data (Fig. 3). We first examined neural patterns across all channels or within frontal, central, or posterior regions of interest. Reward coding was observed in all channels (window tested = 0-3500 ms from reward cue onset), where a cluster in the theta and alpha bands appeared immediately after the reward cue and lasted until shortly before the task cue onset (cluster window = 93-1007 ms, 5.5-15 Hz, corrected p<0.0001). A second, slightly lower-frequency, cluster occurred after the target onset (cluster window = 2253-3007 ms, 2-13 Hz, p<0.0001). In the frontal ROI, a cluster in the alpha band could be observed after the reward cue and lasted until after the onset of the task cue (cluster window = 133-1407 ms, 8.5-14 Hz, p<0.0001). A second cluster in the theta and alpha bands also occurred after the target onset (2253-2967 ms, 2-13 Hz, p=0.0009). There were three clusters in the central ROI: the first one appeared in the high alpha and low beta bands shortly before the task cue and lasted until right before the target cue onset (cluster window = 853-1747 ms, 11.5-15 Hz, p=0.0069). The second cluster also occurred in the alpha and low beta bands after the target cue onset (cluster window = 2373-2907 ms, 9.5-14 Hz). The third cluster was observed between the delta (<4 Hz) and theta bands, also after the target cue onset (cluster window = 2453-3027 ms, 2.5-5 Hz, p=0.0244). Finally, one cluster was found in the posterior ROI in the alpha band after the target cue onset (cluster window = 2513-3027 ms, 8.5-11 Hz, p=0.0423).

**Figure 3.**
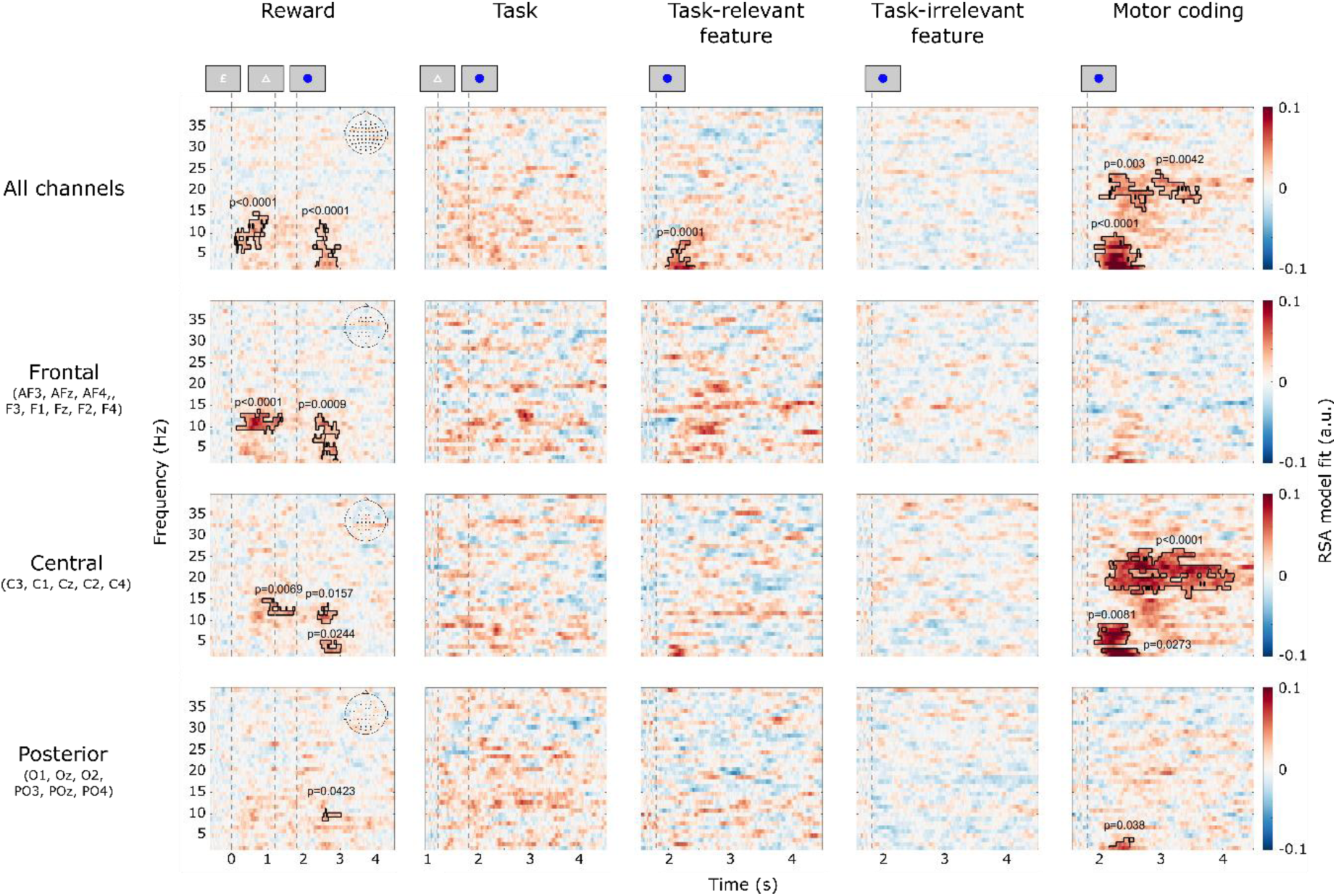
Average neural coding of different task-related features in the time-frequency domain. Each row corresponds to a different ROI: all, frontal, central, or posterior channels; and each column corresponds to a different element: reward level, task type, task-relevant and -irrelevant features, and motor coding. Highlighted areas contain significant clusters at the α=0.05 level.

The target’s task-relevant feature (i.e., blue or yellow on colour trials, and circle or square on shape trials) was only briefly represented in all channels (window tested = 0-2700 ms from the target cue onset), in the low frequencies (cluster window = 153-627 ms, 2-8 Hz, p=0.0001). The task type and the target’s task-irrelevant feature (e.g., blue or yellow on shape trials) were not represented in the time-frequency domain (corrected p>0.05). Finally, the (left-or right-hand) response was represented in all channels (window tested = 0-2700 ms from the target cue onset), with clusters in the delta, theta, and lower alpha bands (cluster window = 113-927 ms, 2-10 Hz, p<0.0001), and in the beta band (cluster windows = 193-1067 ms, 15.5-23.5 Hz, p=0.003; 1053-1847 ms, 17.5-24.5 Hz, p=0.0042). In the central ROI, an alpha cluster occurred immediately after the target cue (cluster window = 113-687 ms, 4.5-9 Hz, p=0.0081). A delta-frequency cluster (cluster window = 233-847 ms, 2-3 Hz, p=0.0273), and a large beta cluster (293-2387 ms, 15.5-26.5 Hz, p<0.0001) were also observed briefly after the target cue.

### Reward prospect desynchronises alpha-beta oscillations

The multivariate analysis indicated that the pattern of alpha-band oscillations differed somehow between reward levels, but does not reveal what the difference was. To better understand how oscillations differed between reward levels, we next used univariate comparisons of the time-frequency response between reward conditions (Fig. 4). Given that many induced effects might dilute when averaging over all channels together in a univariate analysis, we limited the analysis in this section to each individual ROI to capture localized oscillatory changes.

**Figure 4.**
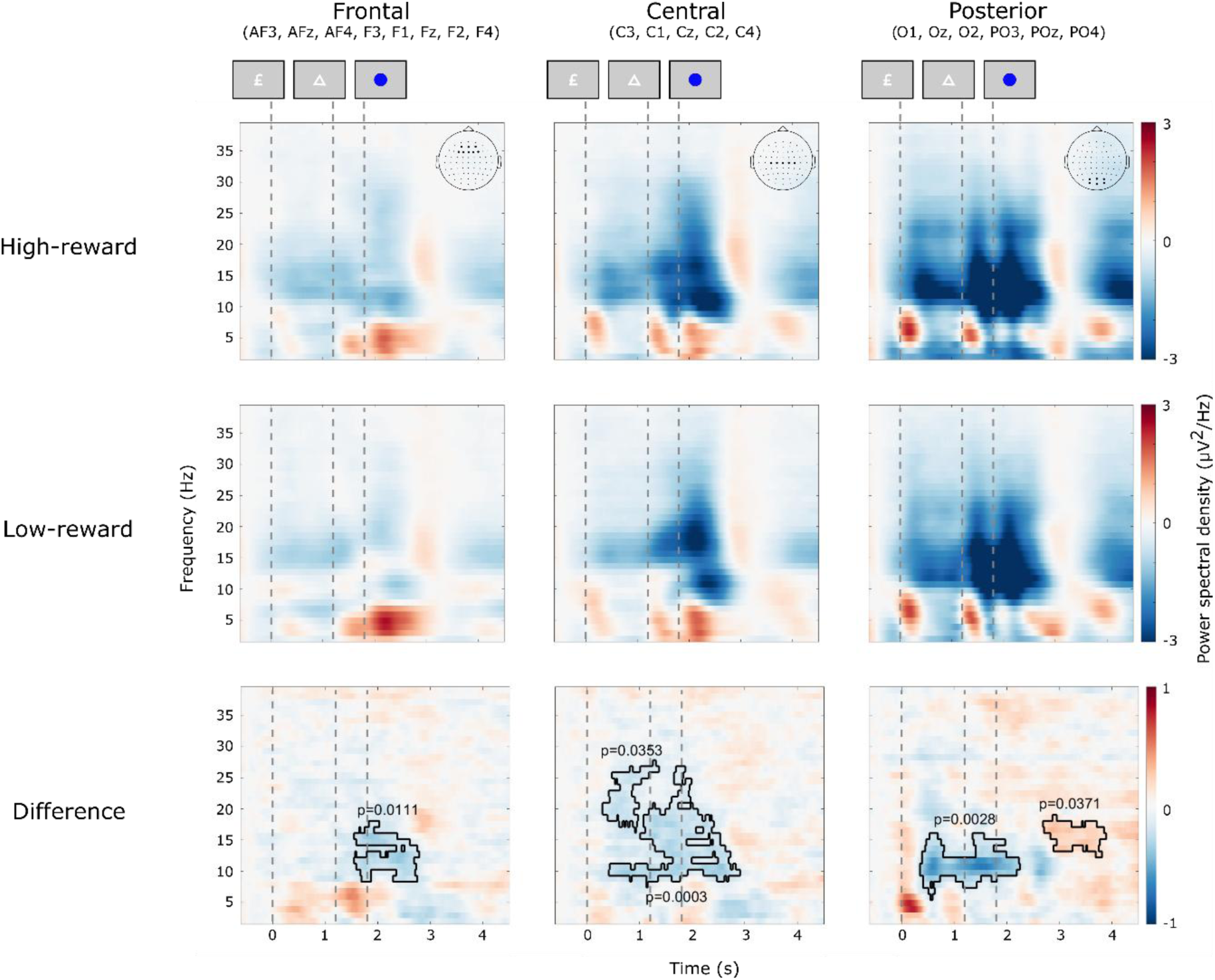
Average power spectral density for different reward conditions. Each row corresponds to a different trial grouping: low-reward, high-reward, and difference between high- and low-reward trials; and each column corresponds to a different ROI: frontal, central, or posterior channels. Highlighted areas contain significant clusters at the α=0.05 level.

Relative to the pre-trial baseline, both alpha and beta bands desynchronised throughout most of the trial, particularly in the central and posterior ROIs. When comparing average power between high- and low-reward trials, a significant cluster was observed in the frontal ROI (window tested = 0-3500ms from reward cue onset) in the alpha and beta bands shortly before the target cue (cluster window = 1553-2787 ms, 8.5-18 Hz, p=0.01). Two larger alpha and beta band clusters appeared in the central ROI, after the reward cue onset, and lasted until after the target cue (cluster windows = 293-1362 ms, 16.5-27.5 Hz, p=0.038; 413-2907 ms, 7.5-26.5 Hz, p<0.0001). A strong desynchronisation in the alpha and lower beta bands was observed in the posterior ROI, starting during the reward phase, and ending after the target cue onset (cluster window = 353-2242 ms, 5.5-16 Hz, p=0.003). Finally, a beta synchronisation cluster appeared in the same ROI by the end of the trial (cluster window = 2693-3887 ms, 12.5-19 Hz, p=0.044). Therefore, oscillatory responses to reward prospect manifested in widespread desynchronisation starting at posterior sites in the alpha band spreading to more anterior sites and higher frequencies around the time of the task cue presentation and subsequent task execution.

### Reward effects on behaviour predict reward sensitivity of alpha-beta modulation

We then subdivided our sample based on a median split of the RT differences between high- and low-reward trials. Participants with larger RT differences unsurprisingly had significantly lower RTs when reward was higher (t(14)=7.08, p<0.0001) but this difference was only at a trend level in the other half (t(14)=2.02, p=0.0624).

Moreover, participants with a larger RT benefit on high-reward trials presented significant reward-related desynchronisation in alpha and beta power in the frontal (window tested = 0-3500 from reward cue onset, cluster windows = 313-2367 ms, 20-26.5 Hz, p=0.0036; 873-2827 ms, 7.5-14 Hz, p=0.0227), central (cluster windows = 253-2327 ms, 4.5-22 Hz, p=0.0023; 453-1647 ms, 19-27.5 Hz, p=0.0331), and posterior (cluster window = 313-2227 ms, 5.5-14 Hz, p=0.0049) ROIs (Fig. 5A). By contrast, participants with lower RT differences between reward conditions showed no significant oscillatory modulation (Fig. 5B).

**Figure 5.**
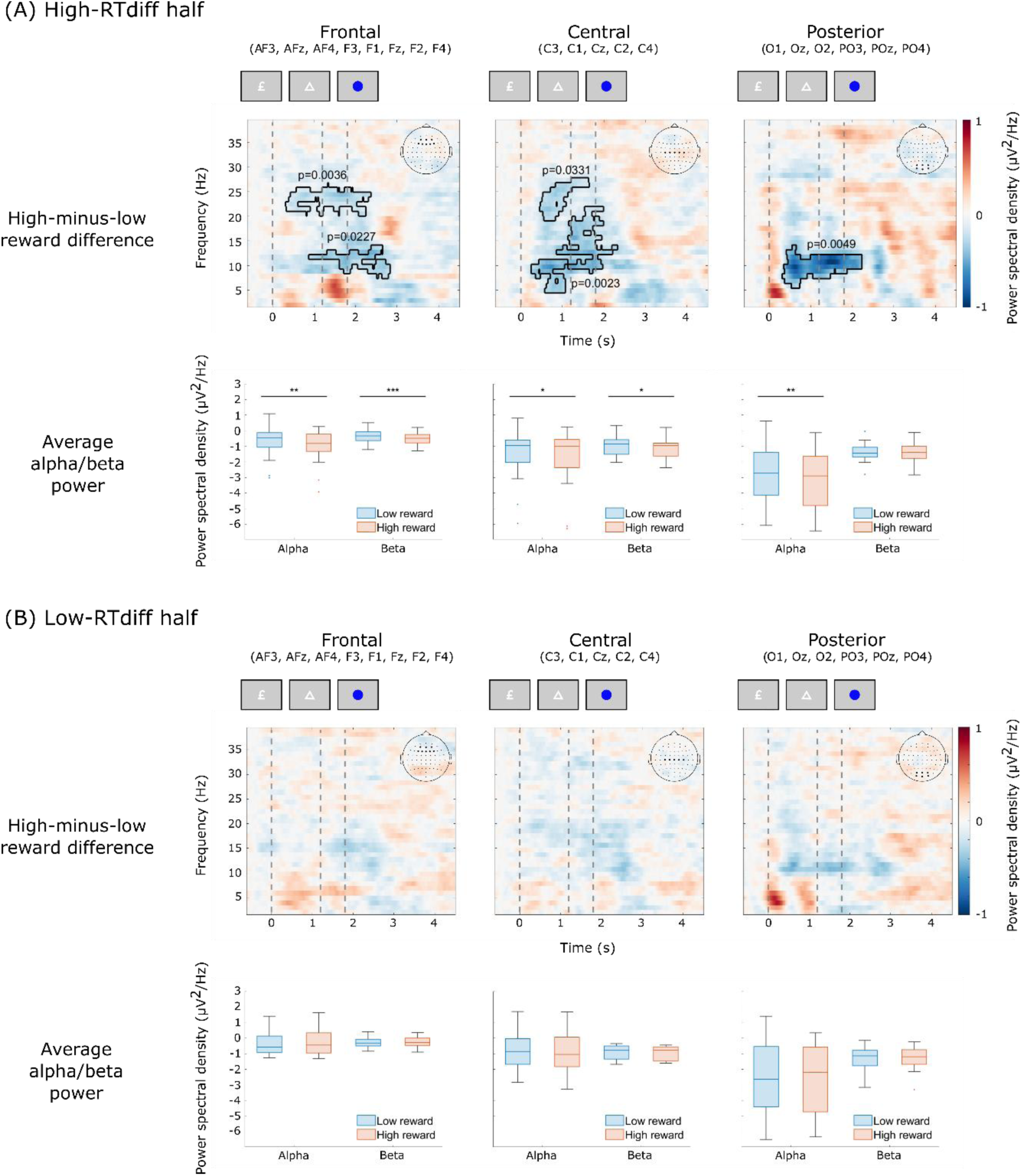
Average power spectral density for participants split by their RT differences in low- and high-reward trials. The first row corresponds to the average difference between high- and low-reward trials, and the second row shows the average alpha/beta power for low (blue) and high (red) reward trials; and each column corresponds to a different ROI: frontal, central, or posterior channels. (A) Participants who had a higher RT difference between reward conditions. (B) Participants who had a lower RT difference between reward conditions. Highlighted areas contain significant clusters at the α=0.05 level. Significance level for pairwise comparisons * = 0.05, ** = 0.01, *** = 0.001.

We also performed individual t-tests on the alpha (window tested = 500-2500 ms from the reward cue onset, 8-12 Hz) and beta (window tested = 500-2500 ms from the reward cue onset, 20-30 Hz) power bands to confirm significant differences across reward conditions for each half of the split. For the high-RT-difference group, we found a stronger desynchronisation difference in the frontal ROI for both alpha (t(14)=3.45, p=0.0039) and beta (t(14)=4.87, p=0.002) power. Similar results were observed in the central ROI for alpha (t(14)=2.9199, p=0.0112) and beta (t(14)=2.58, p=0.0217) power. However, in the posterior ROI, the desynchronisation was only stronger in alpha (t(14)=3.75, p=0.0022) and not in beta power (t(14)=-0.35, p=0.729). Conversely, for the low-RT-difference group, there were no desynchronisation differences in the frontal ROI for either alpha (t(14)=0.39, p=0.6994) or beta (t(14)=-0.73, p=0.4791) power. Similarly, no desynchronisation was found in the central ROI for alpha (t(14)=1.17, p=0.2613) or beta (t(14)=1.23, p=0.2379) power. Finally, no desynchronisation was observed in the posterior ROI for alpha (t(14)=1.77, p=0.0978) or beta (t(14)=0.11, p=0.9135) power.

### Individual differences in alpha and beta power predict response times differences between reward levels

While the above analysis showed a strikingly different pattern of results between groups, we did not test whether the groups themselves differed significantly. To further establish the role of oscillatory power during the reward phase and then compare the relative contribution of different oscillations to performance, we next used signed-rank Spearman correlations to confirm that higher alpha power differences translated into a higher RT difference between reward levels across participants. Average alpha power differences (averaged over 500-1000 ms from the reward cue onset and between 8-12 Hz) correlated with RT differences over all channels (ρ(28)=0.3651, p=0.048), as well as the posterior (ρ(28)=0.3628, p=0.0495) ROIs, and there was a trend in the frontal (ρ(28)=0.3571, p=0.0534) ROI. No significant clusters were found in the central (ρ(28)=0.2988, p=0.1089) ROI.

We then used a cluster-based permutation signed-rank correlation test to find time-frequency windows that predicted the RT difference across reward levels (Fig. 6). This revealed two significant clusters in all channels along the trial (window tested = 0-3500 ms from the reward cue onset): one in the alpha band (cluster window = 193-2367 ms, 6.5-14 Hz, p=0.0091), and one in the beta band (cluster window = 353-2067 ms, 22-29.5 Hz, p=0.0118). The beta cluster could also be found in the frontal ROI (cluster window = 353-2647 ms, 22-26 Hz, p=0.0025), while the alpha cluster was observed in the posterior ROI (cluster window = 373-1507 ms, 3.5-11 Hz, p=0.0091).

**Figure 6.**
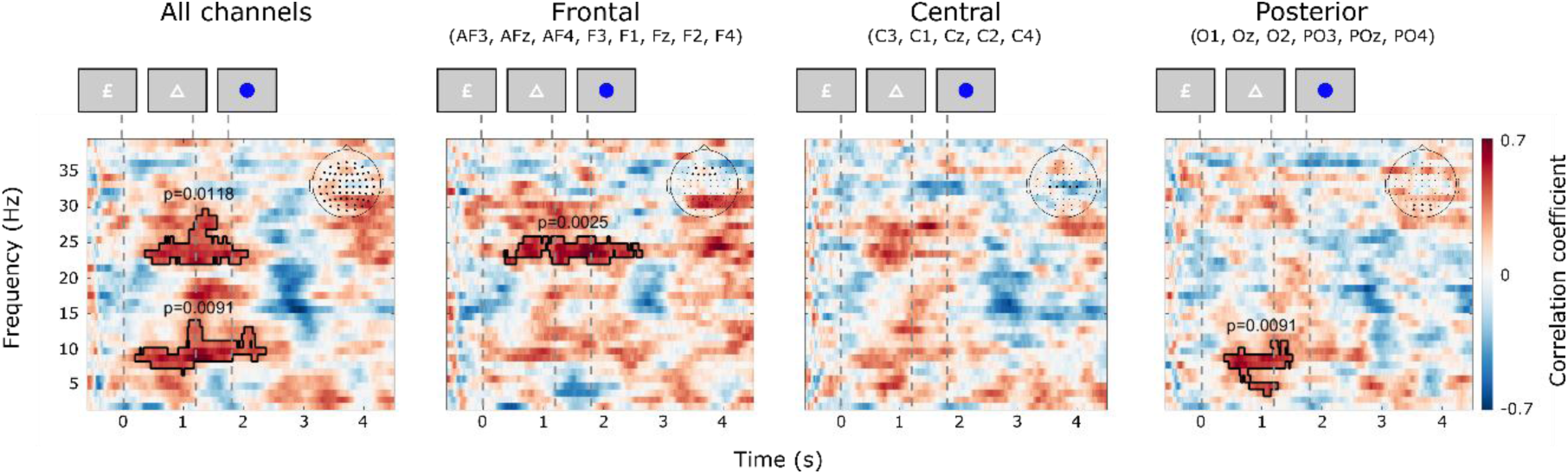
Correlation between desynchronisation differences and response time differences for low and high reward trials. Each plot corresponds to a different ROI: all, frontal, central, and posterior channels, respectively.

### Neural desynchronisation makes a distinct contribution beyond ERP-based neural coding

Finally, we investigated whether the reward-related information in the broadband ERP signal and the oscillatory components observed in the alpha and beta band made independent contributions to the behavioural response. We replicated the RSA analysis of ERP data from Hall-McMaster et al. (2019; Fig. 7A), which found significant neural coding of reward information throughout the trial (cluster window = 67-2973 ms, p=0.0002). We then calculated partial correlations between the average alpha power differences (averaged over 0-1000 ms from the reward cue onset, between 8-12 Hz), the average neural coding (0-1000 ms from the reward cue onset), and the difference in median RTs (RT_diff_) between reward levels (Fig. 7C).

**Figure 7.**
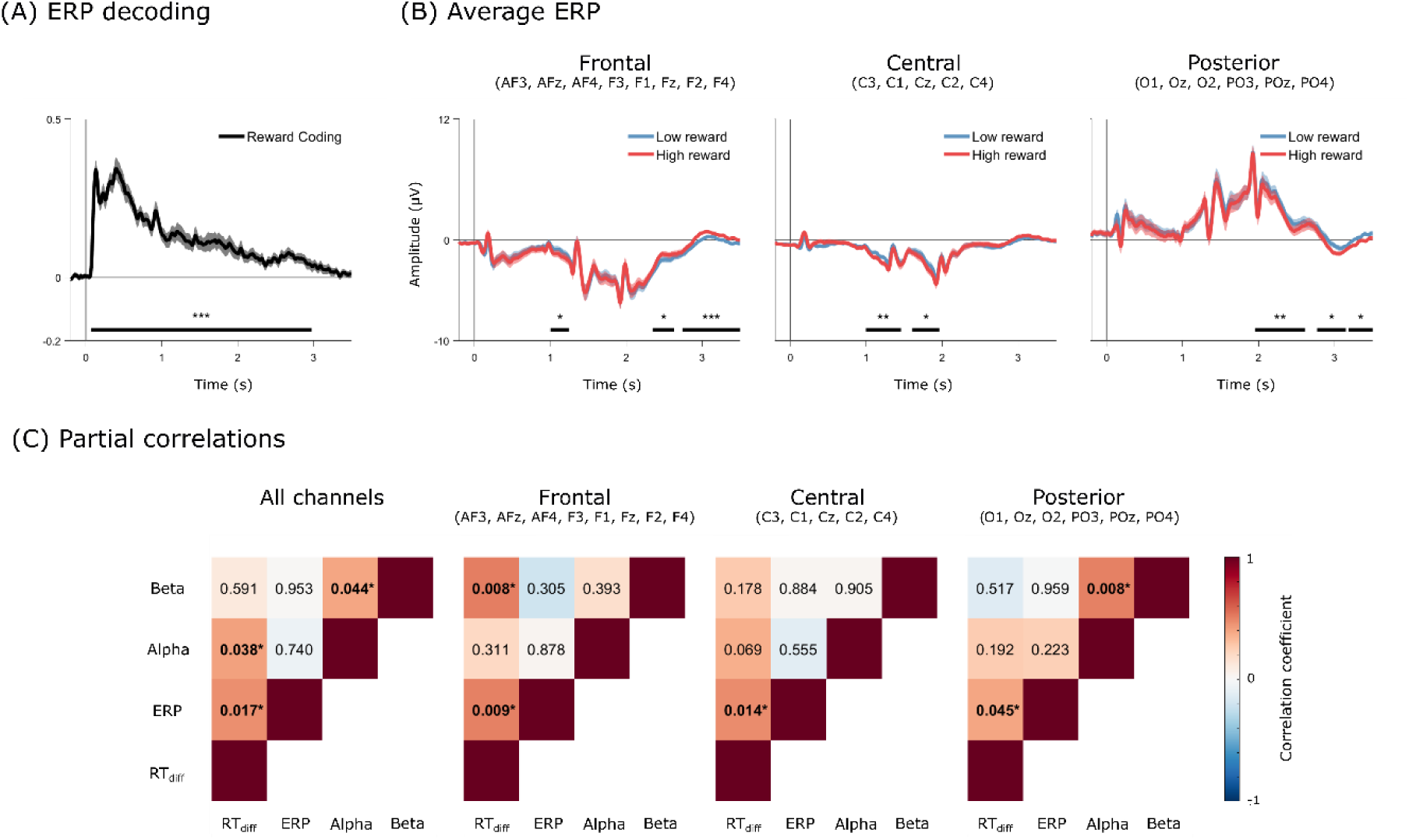
ERPs and contributions from different band powers and the ERP neural coding to RT. (A) Average ERP model fit replicated from Hall-McMaster et al. (2019). (B) Average low- and high-reward ERPs for the frontal, central, and posterior ROIs. The black horizontal bars indicate significant clusters at the * = 0.05, ** = 0.01, *** = 0.001 significance levels. (C) Partial correlations between alpha power, beta power, ERP model fit, and RTdiff across reward levels. Colours denote the correlation coefficient, and numbers indicate p-values.

To explore whether reward cues elicited differences in ERP amplitude, we conducted a cluster-based permutation test comparing high- and low-reward trials across frontal, central, and posterior ROIs (Fig. 7B). We found significant ERP differences (window tested = 0-3500 ms from the reward cue onset) in the frontal (995-1245 ms, p=0.047; 2339-2629 ms, p=0.0186; 2763-3293 ms, p=0.0048), central (991-1489 ms, p=0.004; 1607-1961 ms, p=0.014), and posterior (1975-2557 ms, p=0.0094; 3231-3500 ms, p=0.047) ROIs; however, most of the found differences did not belong to the reward phase. Interestingly, these ERP differences appear to be primarily driven by participants in the high-RT_diff_ group, as no significant clusters could be found in the low-RT_diff_ group (see Supplementary Figure S1 for the differences on each half of the median split).

When averaging alpha power across all channels and controlling for age and sex, signed-rank Spearman partial correlations revealed that stronger ERP-based reward neural coding (ρ(25)=0.4665, p=0.014) and stronger alpha desynchronisation (ρ(25)=0.4973, p=0.008) predicted a higher RT_diff_ , but there was no correlation between ERP decoding and alpha desynchronisation (ρ(25)=-0.0692, p=0.732). Therefore, alpha desynchronisation and reward neural coding appear to be independent predictors of performance. Additionally, alpha desynchronisation was significantly correlated with age (ρ(25)=-0.5421, p=0.004), indicating that age may influence the degree of alpha modulation in response to reward.

We next tested whether beta desynchronisation (0-1000 ms from the reward cue onset, 20-30 Hz) independently predicted performance by including alpha and beta power, as well as reward coding in the ERP data and RT_diff_ in a partial correlation analysis. When averaging alpha power or beta power across all channels and controlling for age and sex, signed-rank Spearman partial correlations also revealed that stronger reward coding in the ERP data (ρ(24)=0.4623, p=0.017) and alpha desynchronisation (ρ(24)=0.4095, p=0.038) predicted a higher RT_diff_, and that the alpha and beta desynchronisation were correlated (ρ(24)=0.3979, p=0.044). However, beta desynchronisation (ρ(24)=0.1104, p=0.591) did not predict RT_diff_, possibly because it competed with alpha power for the same variance since they were themselves correlated. When we restricted alpha and beta power to channels from the frontal ROI, ERP reward coding still predicted higher RT_diff_ (ρ(24)=0.5017, p=0.009), but now RT_diff_ was separately predicted by stronger beta desynchronisation (ρ(24)=0.5107, p=0.008), with no significant correlations between alpha desynchronisation and any of the other measures (p>0.05). Stronger ERP reward coding also led to higher RT_diff_ in the central (ρ(24)=0.4765, p=0.0139) and posterior (ρ(24)=0.3966, p=0.045) ROIs. Interestingly, after taking ERP reward coding into account, alpha or beta desynchronisation measured in the central or posterior ROIs no longer correlated with RT_diff_ (central alpha: ρ(24)=0.3621, p=0.069; central beta: ρ(24)=0.2724, p=0.178; posterior alpha: ρ(24)=0.2643, p=0.192; posterior beta: ρ(24)=-0.1332, p=0.517). Alpha and beta desynchronisation were also correlated (ρ(24)=0.5064, p=0.008) in the posterior ROI. Overall, while the reward coding seems to be largely independent of both alpha and beta desynchronisation and accounting separately for performance improvements, oscillations in the two bands tended to be correlated and therefore accounting for similar variance in RT_diff_.

## Discussion

This study aimed to investigate how reward modulates neural oscillations, and their effects on behavioural performance. We used time-frequency RSA to explore changes in the neural coding of task-related variables, finding significant representations of reward magnitude and motor coding in the spatial pattern of low-frequency oscillatory power. We next looked at desynchronisation changes in different frequency bands as a function of reward, finding that anticipation of larger rewards led to greater alpha and beta desynchronisation. Moreover, across participants the difference in alpha and beta desynchronisation between high and low reward was correlated with differences in RT across reward levels, corroborating the behavioural relevance of reward-related oscillatory changes. Finally, we observed that differences in ERP reward coding and oscillatory desynchronisation between reward magnitude conditions were both predictive of reward-induced performance benefits, suggesting independent mechanisms.

We observed that the representation of reward magnitude, features, and responses were distinct in the time-frequency domain. Reward coding was prominent across all channels and specific regions of interest (ROIs), indicating that reward cues are robustly encoded in the brain’s oscillatory activity. Etzel et al. (2016) had previously evidenced that reward cues enhanced the discriminability of task representations in frontoparietal BOLD signals. This study replicates these results using EEG recordings, while also providing evidence of both alpha and beta oscillations being involved in this process. The presence of reward-related oscillations in the alpha and beta bands suggests that these frequencies play a crucial role in encoding motivational signals. An interesting difference with the results found in Hall-McMaster et al. (2019) is that the task type could not be decoded in the time-frequency domain. Two possible explanations for this could be that the neural patterns associated with this specific task feature are either only phase-locked, or the main components of the elicited ERPs are contained in lower frequencies that fell outside our analysed frequency range (e.g., below 1 Hz). These possibilities could be empirically distinguished by separating the phase-locked and non-phase-locked components of the signal (Singhal et al., 2023), or by applying a high-frequency filter before attempting to decode task-related signals using RSA.

Our analysis revealed significant desynchronisation in the alpha and beta bands across different ROIs in high-reward trials. The alpha modulation is consistent with the idea of an increase of allocated attentional resources (Klimesch, 2012; Palva & Palva, 2007) and the subsequent enhancement of task-relevant processing (Foxe & Snyder, 2011; Heuer et al., 2017; Jensen & Mazaheri, 2010) as a function of reward prospect. While the changes in the beta band were unexpected, we believe they could also be related to reward expectation and task cost anticipation (Gheza et al., 2018), task rule retrieval from long-term memory (Hanslmayr et al., 2012), changes in spatial attention (Sauseng et al., 2005; Siegel et al., 2008), or to anticipated visuomotor planning (Kilavik et al., 2013).

We found that participants with higher RT differences between reward conditions exhibited significant desynchronisation changes in alpha and beta power, whereas those with lower RT differences did not show such modulation. This suggests that individual differences in alpha power modulation may underlie variations in cognitive flexibility and performance. Our results are also consistent with Sawaki et al. (2015), who provided evidence of alpha desynchronisation being associated with faster RTs in high-reward conditions. Here, we additionally found that beta-band desynchronisation may also enhance behaviour. These individual differences in reward processing may be related to the participants’ intrinsic motivation within the task, as those who are more intrinsically motivated might engage more robustly with the reward cues, leading to greater neural modulation and improved performance (Lee & Reeve, 2017). This variability may also reflect broader motivational traits, as suggested by prior work linking reduced alpha modulation to diminished reward responsiveness in depression (Messerotti Benvenuti et al., 2019) and apathy in Parkinson’s disease (Zhu et al., 2019). Such findings support the idea that individual differences in neural sensitivity to reward may stem from underlying differences in motivational engagement.

The correlational analysis revealed that higher alpha power differences were associated with greater RT differences across reward levels. This relationship was particularly strong in the posterior ROI, between the reward and target cues, and may reflect a shift in spatial attention through an increase in cortical sensitivity towards the upcoming target cue (Klimesch, 2012; Rihs et al., 2009). Beta power differences in the frontal ROI, on the other hand, may indicate enhanced readiness and efficiency in executing the task, driven by the motivational influence of anticipated rewards (Zaepffel et al., 2013). Finally, we found that the ERP reward coding through RSA and alpha/beta desynchronisation differences provided individual contributions to the RT differences across reward levels. We believe that the ERP contribution could be a representation of motivation as a function of reward level, as had already been highlighted by Hall-McMaster et al. (2019), while the oscillatory changes may represent a distinct attentional component, showcasing the relevance of exploring multidimensional EEG recording analysis methods to expand on the interaction between ERP and oscillatory components. Interestingly, the partial correlation analysis did not reveal a significant contribution of alpha desynchronisation to RT differences in the posterior region. This could be due to the chosen time window size being larger than the duration of the desynchronisation.

Additionally, we observed a significant correlation between alpha desynchronisation and age, suggesting that individual differences in reward-related alpha modulation may decline with age, possibly due to reductions in attentional engagement or cortical excitability. This aligns with previous findings showing that alpha oscillatory dynamics change across the lifespan, often reflecting shifts in cognitive control efficiency and neural inhibition mechanisms (Voytek et al., 2015).

In conclusion, our study highlights the importance of alpha and beta oscillations in the neural coding of task rules and their modulation by reward. Our results provide evidence for reward directly influencing alpha/beta desynchronisation, particularly during the reward cue phase of the task, which then translates into a faster RT. Finally, we found that ERP reward coding and alpha/beta modulation contribute independently to behaviour, possibly suggesting that the modulation in the alpha and beta bands might not necessarily be related to a representation of reward, but to an increased engagement with proactive control.

## Declaration of Generative AI and AI-assisted technologies

During the preparation of this work the authors used Microsoft Copilot as a peer reviewer and to check for consistency/typographical errors. After using this tool/service, the authors reviewed and edited the content as needed and take full responsibility for the content of the publication.

## Author Contributions

**Juan M. Chau:** Conceptualization, Methodology, Software, Validation, Formal analysis, Writing – Original Draft. **Matias J. Ison:** Conceptualization, Writing – Review and Editing, Supervision. **Paul S. Muhle-Karbe:** Conceptualization, Investigation, Writing – Review and Editing. **Mark. G. Stokes:** Conceptualization, Investigation. **Sam Hall-McMaster:** Conceptualization, Formal analysis, Investigation, Writing – Review and Editing. **Nicholas E. Myers:** Conceptualization, Methodology, Investigation, Writing – Review and Editing, Supervision.

## Supplementary Material

**Table S1.**
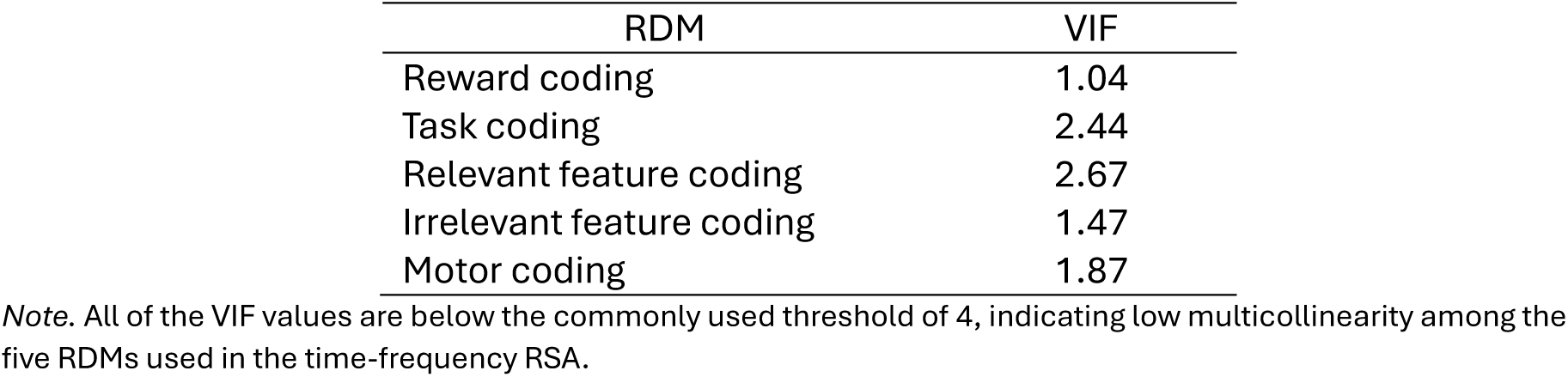
Variance Inflation Factor (VIF) values for the five Representational Dissimilarity Matrices (RDM) used in the time-frequency RSA

**Figure S1.**
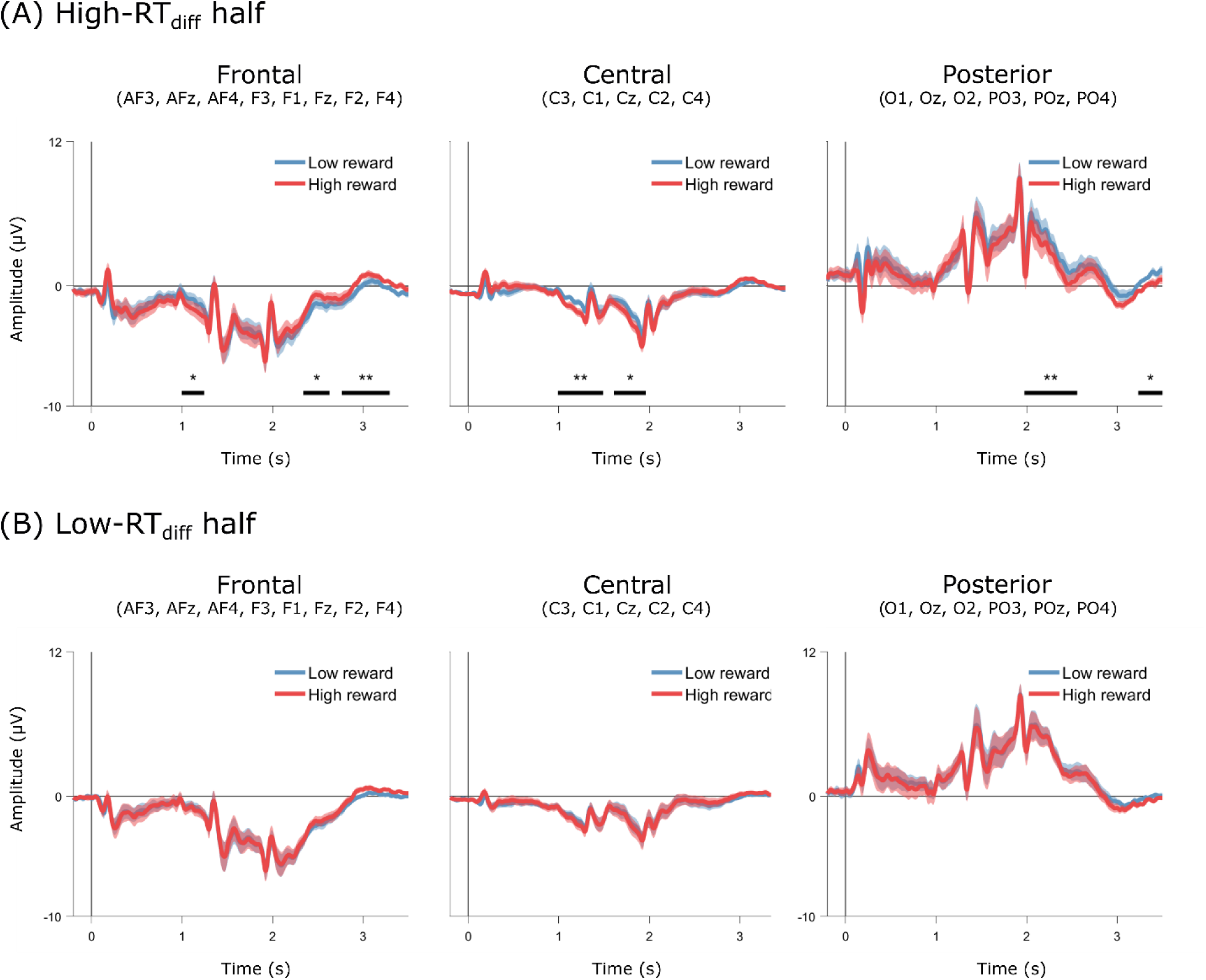
Average high- and low-reward ERPs across the frontal, central, and posterior ROIs. (A) Average ERPs for participants on the high RTdiff half of the split. (B) Average ERPs for participants on the low-RTdiff half of the split. The black horizontal bars indicate significant clusters at the * = 0.05, ** = 0.01, *** = 0.001 significance levels.

